# Neural dynamics of relational memory retrieval across eye movements

**DOI:** 10.1101/2025.03.27.645308

**Authors:** Andrey R. Nikolaev, Roger Johansson, Inês Bramão, Mikael Johansson

## Abstract

Relational memory retrieval entails a dynamic interplay between eye movements and neural activity, yet the temporal coordination of these processes remains unclear. This study examines the relationship between theta and alpha EEG activity and sequential fixations during relational memory retrieval. Participants completed a two-alternative forced-choice associative memory task while their eye movements and EEG were recorded simultaneously. Results reveal a relational eye-movement effect: correctly remembered target elements were disproportionately fixated during later stages of the retrieval sequence. Time-frequency EEG analyses across the test trial show that successful retrieval was characterized by an early, transient increase in theta power and a sustained decrease in alpha power. Gaze-related analyses linked these effects to distinct retrieval processes. The theta effect emerged after the initial cue fixation and predicted retrieval success independently of whether subsequent saccades were directed to the target or distractor. In contrast, the alpha decrease predicted retrieval success only across continued target fixations. These findings connect the relational eye-movement effect to two distinct neural signatures: a nonspecific theta increase, reflecting recollection that may initiate pattern completion, and a target-selective alpha decrease, reflecting sustained reactivation of goal-relevant associations across fixations. Together, these results clarify the temporal dynamics of relational memory retrieval and underscore the role of sequential eye movements in memory-guided behavior.

## Introduction

Eye movements are closely linked to memory for relationships between different elements within an experience (Ryan, Shen, & Liu, 2020; Ryan & Villate, 2009; Smith, Hopkins, & Squire, 2006). Beyond gathering visual information, eye movements actively contribute to relational memory formation by integrating multiple aspects of an experience into a coherent whole (Johansson, Nystrom, Dewhurst, & Johansson, 2022; Kragel & Voss, 2022; Nikolaev, Bramao, Johansson, & Johansson, 2023; Olsen et al., 2015; Wynn, Olsen, Binns, Buchsbaum, & Ryan, 2018). By establishing spatial and temporal relationships, eye movements guide the selection and integration of relevant elements, ultimately facilitating the creation of cohesive memory traces that enable effective navigation and interaction with the environment (Kragel & Voss, 2022; Ryan et al., 2020; Voss, Bridge, Cohen, & Walker, 2017).

Extending this understanding, research on memory retrieval has demonstrated that gaze patterns provide direct insights into how relational memories are reactivated during remembering. Eye-tracking studies show that, well before explicitly reporting their memory judgment, participants disproportionately direct their gaze toward elements matching a cue from a previously studied event (Chua, Hannula, & Ranganath, 2012; Hannula & Ranganath, 2009; Hannula, Ryan, Tranel, & Cohen, 2007; Smith & Squire, 2018; Urgolites, Smith, & Squire, 2018). This gaze bias emerges specifically when participants are tasked with identifying a matching element from an array of distractors (cf., Hannula et al., 2010). This sensitivity of eye movements to relational memory is known as the *relational eye-movement effect*. At the neural level, this effect is linked to activation of the hippocampus (Hannula & Ranganath, 2009). These relational memory effects and their associated hippocampal activity have been observed even when participants fail to consciously identify the matching element (Hannula, Baym, Warren, & Cohen, 2012; Hannula & Ranganath, 2009), suggesting that eye movements might reflect relational memories even in the absence of conscious access (Kumaran & Wagner, 2009; Ryals, O’Neil, Mesulam, Weintraub, & Voss, 2019). Together, these findings indicate that memory for relationships between goal-relevant information guides eye movements.

While these findings offer valuable insights into the relationship between eye movements, memory, and neural activity, they also reveal critical gaps in understanding how relational memory retrieval, eye movements, and neural activity are interconnected during the retrieval process. Addressing these questions requires studying the relational eye-movement effect alongside its neural correlates with high temporal resolution. This enables precise tracking of the dynamic interplay between gaze behavior and neural evidence for memory reactivation.

Building on recent findings that unrestricted eye movements during relational memory formation are linked with EEG activity in the theta (4-8 Hz) and alpha (8-13 Hz) frequency bands when interrelated event elements are sampled across fixations (Nikolaev, Bramao, et al., 2023; Popov & Staudigl, 2023), the present study aims to extend these insights to the retrieval of such relations. In episodic memory research, increases in theta power have been observed to support the retrieval of discrete memories across space and time (Herweg, Solomon, & Kahana, 2020; Staresina & Wimber, 2019). Theta oscillations coordinate sequential inputs of information (Herweg et al., 2020) and supports the successful formation of associative memories (Kota, Rugg, & Lega, 2020). Recent research suggests that theta oscillations guide eye movements during memory formation (Kragel, Schuele, VanHaerents, Rosenow, & Voss, 2021; Kragel et al., 2020) and enable the binding of separate event elements into coherent memory representations (Nikolaev, Bramao, et al., 2023). An increase in theta power is often accompanied by a decrease in alpha/beta power, associated with spread cortical engagement and the broad reactivation of memory content (Martín-Buro, Wimber, Henson, & Staresina, 2020; Michelmann, Bowman, & Hanslmayr, 2016). The changes in theta and alpha activity are thought to play essential but distinct roles during both memory encoding and retrieval (Hanslmayr, Staresina, & Bowman, 2016).

To investigate the dynamics of gaze behavior and brain activity during retrieval, we simultaneously recorded participants’ eye movements and EEG while they performed a two-alternative forced-choice associative memory test. In this task, participants were presented with a cue and asked to select one of two alternatives that had been previously studied alongside it during the study phase, which involved a series of multi-element events. This design allowed us to examine how increases in theta power, presumably involved in retrieving relational memory representations, and decreases in alpha power, likely responsible for reactivating memory content (i.e., event elements), manifest during relational memory retrieval across consecutive fixations. In doing so, we were able to map out the temporal dynamics of the relational eye-movement effect and its underlying neural processes on a fixation-by-fixation basis.

Our approach diverges from prior research on the relational eye-movement effect (Chua et al., 2012; Hannula & Ranganath, 2009; Hannula et al., 2007; Smith & Squire, 2018; Urgolites et al., 2018) in two critical ways. First, most prior studies used paradigms where the cue in the memory test had been studied alongside a single associate, allowing for reactivation of the cue-target association already during cue processing. This makes it difficult to evaluate the specific contributions of target and distractor fixations during retrieval. To address this, we introduced multi-element events during the study phase, reducing the likelihood of retrieving a specific associate solely through cue fixations. Second, previous studies typically presented the target alongside multiple distractors in the memory test to enhance sensitivity to gaze biases favoring the target. In contrast, we adopted a two-alternative forced-choice protocol, presenting only one distractor. This design ensured an equal likelihood of fixating either the target or distractor after the initial cue fixation and provided sufficient gaze-related EEG data for both targets and distractors in subsequent analyses.

We expect the present study protocol to elicit a relational eye-movement effect that reliably differentiates between recalled and forgotten element associations. Specifically, this effect is anticipated to align with an increase in theta power accompanied by a decrease in alpha power. By examining this interplay across successive fixations, we aim to uncover how the neural dynamics of the relational eye-movement effect evolve over time.

## Method

### Participants

Thirty-six healthy adults participated in the experiment in exchange for a gift card in a shopping mall (approx. €20). Three participants were excluded due to technical problems, three due to below-chance memory performance, and two due to ceiling memory performance (i.e., too few incorrect trials for the planned analyses). Thus, data from 28 participants (20 females; mean age 23.3 years; age range 18-32 years) were included in the analysis. The study was conducted in accordance with the Swedish Act concerning the Ethical Review of Research involving humans (2003:460). All participants gave written informed consent, and the study followed the local ethical guidelines at Lund University.

### Stimuli and procedure

Stimuli during study and test encompassed a total of 324 grayscale photos of faces (Chelnokova et al., 2014), places (http://www.tarrlab.org), and objects (Brodeur, Dionne-Dostie, Montreuil, & Lepage, 2010) in the shape of a square measuring 4.2 degrees of visual angle (dva) (Fig. 1). All image exemplars used in the experiment were unique and were randomly assigned to events and participants.

**Fig. 1.**
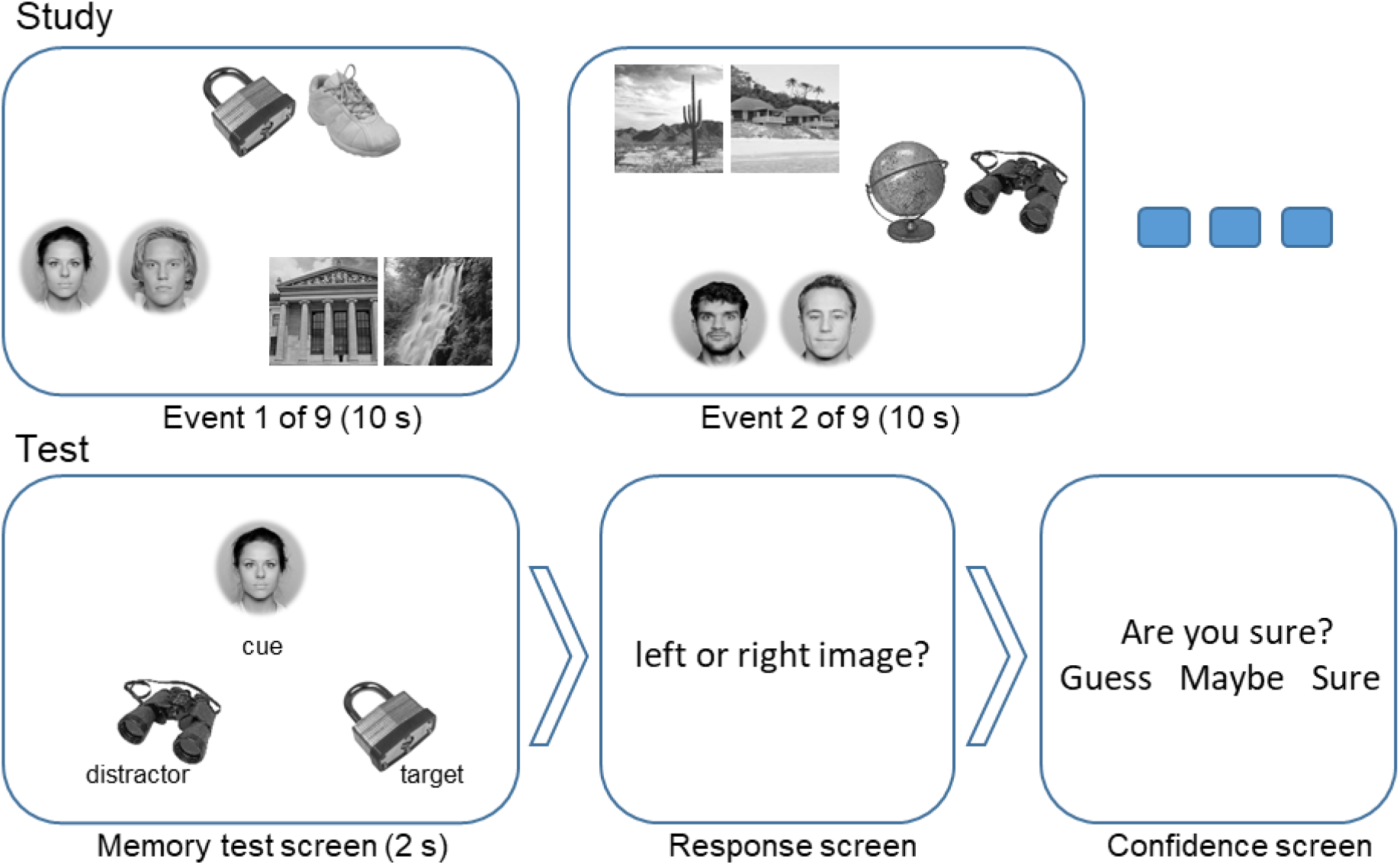
Schematics of the study and test phases of the experiment. During study, nine event screens were presented sequentially for 10 s each. Each event screen comprised six images arranged in three pairs of two exemplars from the same category. The pairs were located along the perimeter of an invisible circle with a radius of 9.2 dva from the screen center. The locations of the pairs on the perimeter were random, but equidistant. In the subsequent memory test phase, test screens were presented for 2 s, comprising three elements: cue, target, distractor. The distance between the cue and the other two elements was 6.5 dva. Participants had to indicate which of the bottom elements (a lock in this example) was from the same event screen as the cue during study. The response was given during the response screen, followed by a confidence question. The response and confidence screens were presented until the answer was given. *Note:* Faces were obtained from the Oslo Face Database (Chelnokova et al., 2014). All individuals whose faces appear in this figure provided permission for their faces to be published, as documented in the associated user agreement form.

The experiment included six blocks with short breaks between them. Each block began with a study phase, comprising nine event screens with six elements from three categories (2 faces, 2 places, 2 objects) presented for 10 s each (Fig. 1A). Exemplars from the same category were placed close together, while those from different categories were placed farther apart. These three category pairs were arranged around an invisible, centered circle, where their locations were randomized for each event, with the pairs were always separated by 120° arcs. Participants were to memorize the multiple inter-category associations specific to each event. The test phase began following a distractor task that required participants to count backward in steps of seven from a randomly selected three-digit number for 30 seconds. During the test, participants were prompted to identify which of two event elements had appeared in the same event as the provided cue element.

The test screen consisted of three elements: cue, target, and distractor, and was presented for 2 s (Fig. 1B). While the cue element was always from a different category than the target and distractor elements, the target and distractor elements were always from the same category. The distractor was selected from another event encoded no more than two screens before or after the target event. Because each cue was associated with four potential targets during the study phase, success in the memory test could only occur after the target and/or distractor was fixated. Including a single distractor ensured an equal likelihood of fixating either the target or the distractor following the initial cue fixation. This design also ensured sufficient gaze-related EEG data for both targets and distractors. The position of these three elements was fixed in all test screens: cue element centrally at the top, and target and distractor elements to the right and left at the bottom (Fig. 1B). The spatial configuration of the elements in the test screen was designed to make participants’ eye movement patterns stereotypical to the task, involving initial gaze towards the cue at the top and then shifting their gaze towards the target and distractor elements at the bottom (Fig. 1D). After each test screen, participants were to make their memory judgment (Fig. 1C). Their response was registered by pressing the left or right arrow key (corresponding to the locations of where the target and distractor were presented in the preceding test screen). Finally, they also indicated their choice confidence by selecting “Sure”, “Maybe”, or “Guess” in response to the question “Are you sure?”. Choice confidence was registered by the left, down, and right arrow keys, respectively. The relative target and distractor location (right or left) was randomized for each test trial.

The memory test was designed to assess relational memory for event specific inter-category element associations. To this end, inter-category associations were tested for each element in each event on four trials. For example, a face element was tested twice with place elements (once as cue and once as target), and twice with object elements (once as cue and once as target). The cue-target status of the categories was counterbalanced across blocks so that in the odd blocks the cue-target status was faces-places, places-objects, objects-faces, and in the even blocks the cue-target status was places-faces, objects-places, faces-objects. Thus, 12 different associations were tested for each event. In total, 108 associations were tested in each block (12 × 9 = 108 tests for the nine events within a block) with randomized order of presentation. A fixation cross was shown for a random interval from 1 to 1.5 s before each study and test phase.

Two practice sessions with increasing difficulty preceded the main experiment. The first session comprised three event screens in the study phase, and the second session comprised six event screens. The increased difficulty prepared participants for the main experiment, which comprised nine event screens per study phase. In each practice session, participants completed ten memory tests during the test phase, each followed by feedback on the correctness of their response.

The participant’s head was stabilized with a chin rest at a distance of 62 cm from the monitor. Stimuli were presented on an EIZO FlexScan EV2451 monitor with a resolution of 1920 x 1080 pixels and a refresh rate 60 Hz. Stimulus presentation and behavioral data collection were controlled using PsychoPy v.2020.2.4 (Peirce et al., 2019).

### EEG and eye movement recording

EEG and eye movements were recorded concurrently throughout the experiment. EEG was recorded using a Grael 4 K EEG amplifier (Neuroscan, Compumedics Limited, Australia) at a sampling frequency of 2048 Hz using 31 Ag/AgCl electrodes positioned according to the extended 10-20 International system (Easycap GmbH, Germany). Two electrodes placed over the left and right mastoids were used as the recording reference. The ground electrode was AFz. Additional electrodes recorded the vertical and horizontal electrooculograms (VEOG and HEOG): the VEOG electrodes were placed above and below the right eye, the HEOG electrodes were placed at the left and right outer canthi of two eyes. Electrode impedances were kept below 5 kΩ.

The Tobii Pro Spectrum eye tracker (Tobii, Stockholm, Sweden) recorded movements of both eyes with a sampling frequency of 600 Hz. 9-point calibration and validation routines were conducted before each experimental block. An error of 1 dva between calibration and validation was tolerated: if it was larger, the calibration was repeated. The eye tracking was controlled from the PsychoPy experimental program via the open-source toolbox Titta (Niehorster, Andersson, & Nystrom, 2020). The experimental program ran on the same computer that controlled the eye tracker. To synchronize stimulus presentation and EEG recording, a transistor-transistor logic (TTL) signal was sent via a parallel port from the stimulus presentation computer to the EEG system at the beginning and end of each block, encoding event, and memory test. To ensure synchronization between EEG, eye tracking, and stimulus presentation, we used the Black Box ToolKit (The Black Box ToolKit Ltd., UK), including the optodetector, prior to the experiment. We measured the time differences between the image onset on the screen and the TTL signal, as well as between the image onset on the screen and the stimulus onset marker sent to the eye tracker. Constant differences of 23 and 10 ms, respectively, were identified and corrected by shifting the position of the markers offline.

### EEG analysis

In the present study, only data from the test phase were analyzed (the results of the study phase were reported in Nikolaev, Bramao, et al., 2023). Analysis was conducted with custom scripts for R (version 4.2.0, 2022) and MATLAB R2024a (The MathWorks Inc., Natick, Massachusetts), using the EEGLAB toolbox (Delorme & Makeig, 2004), the EYE-EEG extension (Dimigen, Sommer, Hohlfeld, Jacobs, & Kliegl, 2011) for EEGLAB, and the Unfold toolbox (Ehinger & Dimigen, 2019). Time-frequency analysis and mapping was performed using the FieldTrip toolbox (Oostenveld, Fries, Maris, & Schoffelen, 2011).

To analyze EEG in unrestricted viewing, we employed a previously established processing pipeline (Nikolaev, Ehinger, Meghanathan, & van Leeuwen, 2023), comprising four major steps. First, we synchronized the eye-tracking data with EEG using the EYE-EEG extension for EEGLAB (Dimigen et al., 2011). Second, we applied the set of conventional functions to clean EEG from the EEGLAB toolbox (Delorme & Makeig, 2004). Third, we applied the OPTICAT function, which is specifically designed to remove oculomotor artifacts during unrestricted viewing (Dimigen, 2020). Finally, we adjusted for the overlapping effects of subsequent saccades on the EEG and the confounding effects of oculomotor covariates by applying deconvolution modeling using the Unfold toolbox (Ehinger & Dimigen, 2019). See Nikolaev, Ehinger, et al. (2023) for full details of this approach to EEG preprocessing and deconvolution modeling.

### EEG preprocessing

The EEG preprocessing aimed to clean the EEG obtained during unrestricted viewing. The key operation of the Unfold toolbox for correcting overlapping effects is time expansion, which requires continuous EEG as input. Therefore, all cleaning was performed on the continuous (i.e., non-segmented) EEG by applying the EEGLAB functions as specified below.

The EEG was downsampled to 256 Hz. Power line noise was removed by using multi-tapering and a Thompson F-statistic implemented in the *pop_cleanline* function. Flat-line channels, low-frequency drifts, noisy channels, and short-time bursts were removed using artifact subspace reconstruction (ASR) with parameter 20, implemented in the *clean_artifacts* function (Mullen et al., 2015). Next, oculomotor artifacts were corrected using the ICA-based *OPTICAT* function (Dimigen, 2020), which removes all types of ocular artifacts (e.g., eyeball rotation, blinking), especially focusing on myogenic saccadic spike activity. This function optimized the ICA decomposition by training it on a copy of the high-pass filtered EEG at 2 Hz, in which sampling points between −20 and +10 ms of saccade onset are overweighted. The obtained ICA weights were applied to the unfiltered version of the same EEG data. This function then computed the ratio between the mean variance of the ocular ICA components during saccade and fixation; components exceeding the ratio of 1.1 were removed. The obtained ICA components were also checked for other types of artifacts by using the automatic classifier (*pop_iclabel*) (Pion-Tonachini, Kreutz-Delgado, & Makeig, 2019). On average, 15.9 (SD = 3.2) components were removed per participant. The EEG was re-referenced to average reference. The channels removed during ASR (mean = 1.2, SD = 1.03 per participant) were interpolated using spherical spline interpolation.

We perform three types of theta and alpha power analyses: one at the level of the entire test trial, and two gaze-related analyses: saccade-locked and fixation-locked power analyses. For the analysis at the level of the entire test trial, we first evaluated the time-frequency difference between memory conditions. Specifically, we calculated the power spectrum of 2-s EEG segments for frequencies between 1 and 30 Hz with the multi-taper method using a Hanning taper and a fixed 0.5-second time window for each frequency (*ft_freqanalysis* of the FieldTrip toolbox). The resulting power values were log-transformed, grand averaged, and subtracted (recalled minus forgotten memory test trials). The areas of maximum difference on the plot were used to select the frequency ranges for statistical power analyses: 3-5 Hz for the theta band and 8-13 Hz for the alpha band, as explained in Results.

For statistical analysis of all three types, we used the procedure proposed by (Ossandón, König, & Heed, 2020) to extract theta and alpha power. This procedure yielded continuous power time series suitable for deconvolution for inclusion in the deconvolution model for saccade-locked and fixation-locked analyses. For each analysis type, we processed each EEG channel as follows. First, the cleaned continuous EEG in each channel was band-pass filtered with a zero-phase FIR filter (*pop_eegfiltnew*) in the selected theta and alpha bands. Next, the analytic signal was derived with the Hilbert transform and instantaneous power was computed as the squared modulus. Next, we computed a baseline reference by averaging the power values across all baseline windows pooled over trials using the following analysis-specific baselines. For the entire test trial (0-2000 ms relative to test screen onset), the baseline interval was −200 to 0 ms before test screen onset. For the saccade-locked interval (−200 to +200 ms around saccade onset) the baseline interval was −300 to −200 ms before saccade onset. For the fixation-locked interval (0-400 ms relative to fixation onset), the baseline interval was −200 to −100 ms before fixation onset. Finally, we converted the power values of each trial of interest into decibel changes by taking ten times the base-10 logarithm of the ratio of instantaneous power to mean power in the chosen baseline window. Using analysis-specific baselines converted each segment into relative (dB) measures on the same scale, allowing for direct statistical comparison of EEG power between gaze-related epochs.

Note that that selecting the baseline interval for gaze-related EEG epochs in EEG-eye movement coregistration research in free viewing can be challenging (see Nikolaev, Meghanathan, & van Leeuwen, 2016 for a discussion of this issue). For the current analyses, we used local, event-specific baseline windows that were immediately before each saccade- or fixation-locked interval of interest. We did this instead of using a single, common baseline (e.g., pre-test screen or pre-cue fixation) because our focus was on the retrieval processes that unfolded with eye movements on each screen element. Moreover, ongoing unrestricted eye movements can produce slow drifts and evolving background oscillatory activity across sequential gaze-related EEG epochs (Fischer, Graupner, Velichkovsky, & Pannasch, 2013; Guérin-Dugué et al., 2018; Kamienkowski, Varatharajah, Sigman, & Ison, 2018; Nikolaev, Meghanathan, & van Leeuwen, 2018). Consequently, a distant common baseline becomes progressively less representative of the neural state that is immediately around the time of an eye movement and could artificially amplify or attenuate EEG differences between gaze-related epochs.

### Deconvolution

Correcting for overlapping effects of sequential saccades on the EEG and controlling for the effects of continuous covariates on the EEG, such as oculomotor characteristics (e.g., saccade size, fixation duration) or temporal positions of a fixation (e.g., fixation rank), is an important step in the analysis of EEG co-registered with eye movements in unrestricted viewing (Dimigen & Ehinger, 2021; Dimigen et al., 2011; Nikolaev et al., 2016). To do this, we used the regression-based deconvolution modeling implemented in the Unfold toolbox (Ehinger & Dimigen, 2019). The output of the toolbox was time series of regression coefficients (betas) representing the partial effects of the predictors, adjusted for the covariates included in the model. The time series corresponded to subject-level averaged waveforms in traditional EEG analysis.

The deconvolution modeling for the saccade- and fixation-locked analyses was largely identical. The model included predictors at the level of entire trials and at the level of eye movements. To account for the EEG activity evoked by presentation of the test screen we included in the model its onsets (the intercept term). Similarly, the saccade or fixation onsets were included. The categorical predictors were defined by the combination of the memory outcome (recalled vs. forgotten memory test trials) and the fixated element (cue or target and distractor). Specifically, separate deconvolution models were run for fixations of rank 1 on the cue, fixations of rank 2 on the target and distractor, and fixations of rank 3-4 on the target and distractor. To ensure that the overlapping effect of each fixation of a trial was corrected, fixations not of interest to the current model were accounted for by assigning them to the “other” level of the categorical predictor. We also included fixation rank in the model because it affects gaze-related EEG (Fischer et al., 2013; Guérin-Dugué et al., 2018; Kamienkowski et al., 2018; Nikolaev et al., 2018). The order of memory tests over the course of an experiment (as trial numbers from 1 to 648) was also included, since increasing fatigue and decreasing vigilance during an experiment may affect EEG (Makeig & Jung, 1996; Tran, Craig, Craig, Chai, & Nguyen, 2020). Three eye movement covariates, which are known to affect gaze-related EEG (Cornelissen, Sassenhagen, & Vo, 2019; Dimigen et al., 2011; Nikolaev et al., 2016), were included in the model: saccade size, angle and fixation duration. Fixation duration was included in the model for fixation-locked power but not in the model for saccade-locked power to ensure comparability between the two types of analyses by maintaining exactly the same set of trials and participants. Because eye movement effects on the EEG are nonlinear (Dimigen & Ehinger, 2021; Nikolaev et al., 2016), they were modeled with spline predictors. In general, the model formula was as follows:

~~~
Screen onset: EEG ∼ 1
Fixation or saccade: EEG ∼ 1 + categorical predictor + fixation rank + trial N +
spl(fixation duration,5) + spl(sac_amplitude,5) + circspl(sac_angle,5,-180,180)
~~~

### Statistical analysis

For the EEG analysis of the entire test trial, the dB-normalized instantaneous power in the theta and alpha bands was segmented from −200 to 2000 ms relative to the test screen onset. Power was averaged across trial and baseline corrected by subtracting the mean power in the interval from −200 to 0 ms. The analytical time windows were defined based on the maximal time-frequency differences between recalled and forgotten trials, as described in the Results section.

For the gaze-related EEG analyses, analytical time windows were determined on theoretical grounds, reflecting the temporal dynamics of visual processing and memory retrieval. In the saccade-locked analysis, the window from −200 to 0 ms relative to saccade onset was chosen to capture neural activity associated with preparing eye movements from the cue to the target or distractor, as well as subsequent gaze transitions between elements. In the fixation-locked analysis, the window from 0 to 200 ms after fixation onset was selected to capture neural processes underlying visual information acquisition and encoding during fixation.

The starting time of the fixation-locked window was set at the fixation onset because narrow-band filtering of EEG (theta, alpha) reduces temporal resolution: longer impulse responses introduce temporal smoothing and thus reduce the precision of event-related alignment to saccade or fixation onsets. Additionally, since EEG power was only normalized in dB relative to the reference intervals in the continuous power time series and was not further baseline-corrected after deconvolution, the resulting time windows reflected a combination of event-specific neural responses and ongoing oscillatory activity. Although this approach limited precise event-related temporal accuracy, it allowed us to capture the evolution of retrieval processes across sequential eye movements (see Discussion for further rationale and challenges in selecting analytical windows in free viewing).

We used a priori selected eight groups of three electrodes around the landmark electrodes of the International 10-20 System of Electrode Placement: F3, F4, C3, C4, P3, P4, O1, O2. They comprised eight regions of interest (ROIs) over frontal, central, parietal and occipital brain regions of the left and right hemispheres (FL, FR, CL, CR, PL, PR, OL, OR) (the inset in Fig. S1). EEG was averaged across the three electrodes in each ROI.

Statistical analyses were conducted using repeated-measures ANOVA with the factors of Memory (forgotten vs. recalled), ROI (frontal, central, parietal, occipital), and Hemisphere (left vs. right) on the power values averaged over the time windows. In the analysis of the gaze-related EEG for target and distractor, the factor Element was added to this ANOVA (see below). When presenting EEG results below, we report statistical effects and interactions only when they involve the Memory factor. We used the Huynh-Feldt correction for p-values associated with two or more degrees of freedom to control for sphericity violation and the Tukey test for post-hoc analyses. Statistical analyses were performed using STATISTICA 10 (TIBCO Software Inc., Palo Alto, CA, USA) and R (version 4.4.1, 2024).

## Results

### Recalled and forgotten trials

Test performance was assessed by considering both retrieval correctness (‘correct’, ‘incorrect’) and confidence (‘guess’, ‘maybe’, ‘sure’). These response outcomes thus provide six possible combinations that in theory could be used to disambiguate between different levels of successful relational memory.

The average of correct memory retrieval across participants was 68.6 ± 8.3 % (mean ± SD). The percentage of correct and incorrect responses depending on response confidence is shown in Table 1.

**Table 1.** Percentage of correct and incorrect responses by confidence levels (mean (SD) % across 28 participants)

|  | guess | maybe | sure |
| --- | --- | --- | --- |
| correct | 13 (8.6) | 28 (9.3) | 27 (14.9) |
| incorrect | 11 (6.6) | 16 (8.6) | 4 (3.2) |

To increase the contrast between retrieval outcomes and to maximize the number of trials for the gaze-related EEG analysis, we considered two memory retrieval trials: recalled and forgotten relations. In the recalled trials, we considered test trials that resulted in correct responses with confidence ratings of ‘maybe’ and ‘sure’. In the forgotten trials, we included test trials that resulted in incorrect responses, as well as any trials with a confidence rating of ‘guess’.

On the test screens, the locations of target and distractor images occasionally coincided with their positions at study. Such spatial congruence could, in principle, bias memory performance. To rule out this possibility, we compared accuracy when target images appeared in the same versus the opposite location as during encoding and found no difference between conditions, excluding a systematic effect of spatial congruence. We also examined whether the category of the cue image influenced performance. Accuracy did not differ across faces, places, and objects, ruling out an effect of cue category on memory performance (see Supplementary Materials for details on both controls).

### The relational eye-movement effect

Each fixation within the 2-s memory test trial was given a sequential number, i.e., *fixation rank*, which indicated the visiting order of each element in the screen. The typical eye movement pattern involved first fixating on the cue (fixation rank 1) and then shifting gaze to the elements below, with an equal number of fixations on the target and distractor (fixation rank 2 and subsequent fixations). The number of fixations gradually decreased over the course of a trial (Fig. 2A), so that fixation ranks later than 4 did not provide sufficient data for reliable eye movement and EEG analyses.

**Fig. 2.**
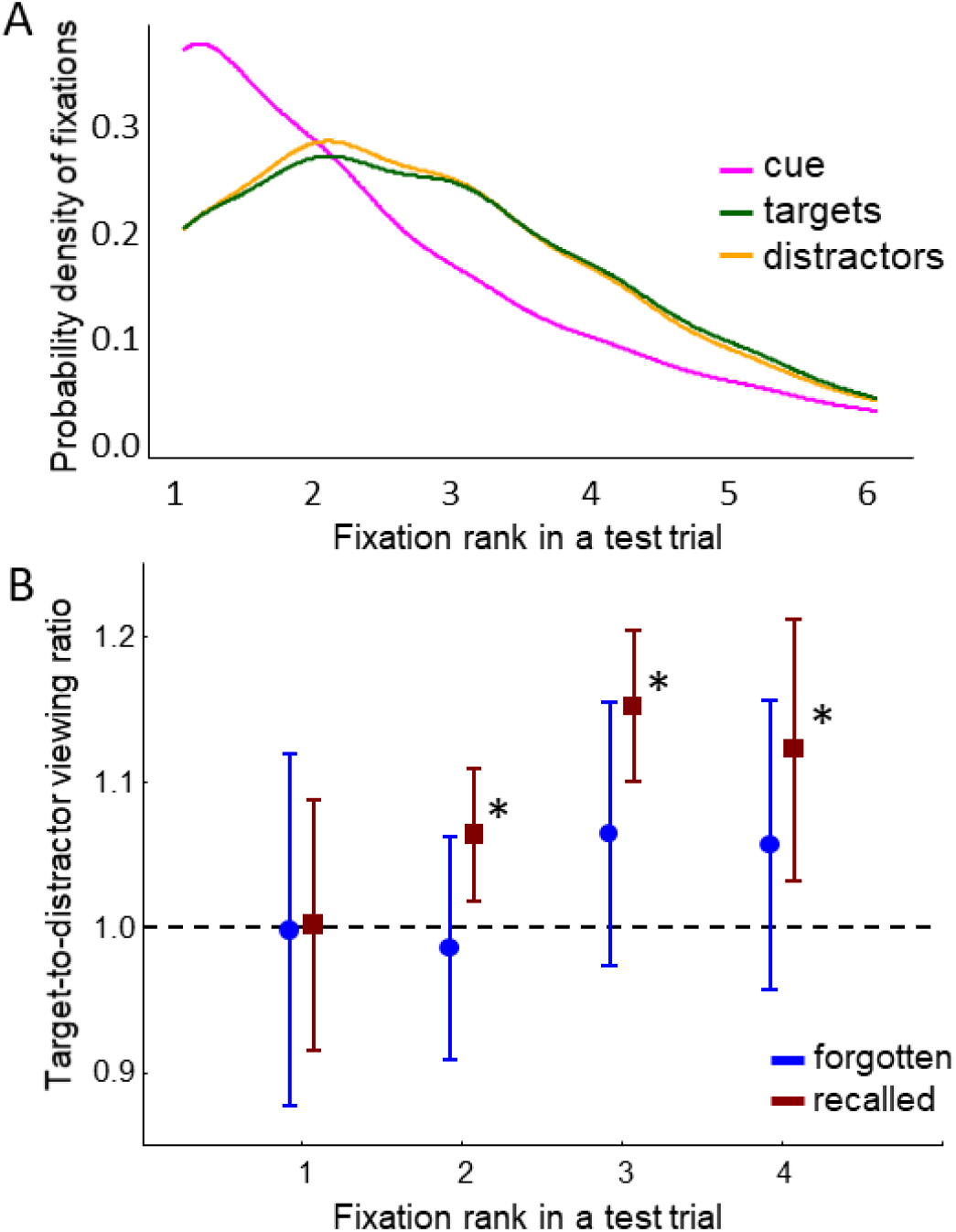
Time course of relational eye movements. (A) Probability density of fixations on cue, target, and distractor during test as a function of fixation rank, i.e., the order of the fixations in a test trial. Thus, in most cases, the test trials started with a fixation on the cue (fixation of rank 1) and continued with fixations on the target or distractor with equal probability (fixations of ranks 2, 3, 4 and so on). Note that continuous curves represent kernel density estimates for visual clarity; fixation rank itself is a discrete variable. (B) Relational eye-movement effect. The target-to-distractor viewing ratio over all participants (N=28) as a function of fixation rank. Values above 1 indicate disproportionate viewing of the target compared to the distractor. Error bars denote 0.95 confidence intervals across participants. Asterisks mark significant differences from 1.

We then verified the presence of the expected relational eye movement effect and examined how it evolved along the fixations rank 1 to 4 with respect to recalled and forgotten relations. To do this, we compared the time spent looking at the target versus the distractor during the 2-s test trial. We computed the duration of fixations on the target and distractor of each rank relative to the mean duration of fixations on all 3 elements within the test trial (similar to Hannula et al., 2007). After we calculated the ratio of these normalized fixation durations between the target and the distractor. A ratio greater than 1 indicates disproportionate viewing of the target compared to the distractor (Fig. 2B).

To examine the time course of target viewing, we compared the target-to-distractor viewing ratios against 1 across fixation ranks using a one-tailed t-test with FDR-corrected p-values. In the recalled, the proportion of viewing time directed to the target was significantly greater than to the distractor for fixation ranks 2-4 (all ps < .02). This effect is further illustrated in Fig. 2B, where the 0.95 confidence intervals for these fixation ranks do not include 1. These findings indicate that the relational eye-movement effect was present for recalled but not for forgotten relations.

### Neural activity during the entire test trial

Figure 3 illustrates the correspondence between successive eye movements and retrieval-related changes in theta and alpha activity over the entire 2-s test trial. The time course of eye movements is shown by the latencies of fixation onsets of the first six ranks, which vary increasingly over time along the retrieval time window (Fig. 3A). The time-frequency plot contrasting recalled and forgotten trials shows that retrieval is associated with a theta increase peaking between 500 and 1000 ms and a gradual alpha decrease starting at 500 ms and lasting until the end of the trial (Fig. 3B). The first fixation, which predominantly occurred on the cue, was not associated with any EEG differences between the two memory conditions. The following fixations, which mainly occurred on target and distractor, at ranks 2 to 4 corresponded to the interval of the higher theta power. In comparison, the fixations at ranks 3 to 6 corresponded to the interval of lower alpha power.

**Fig. 3.**
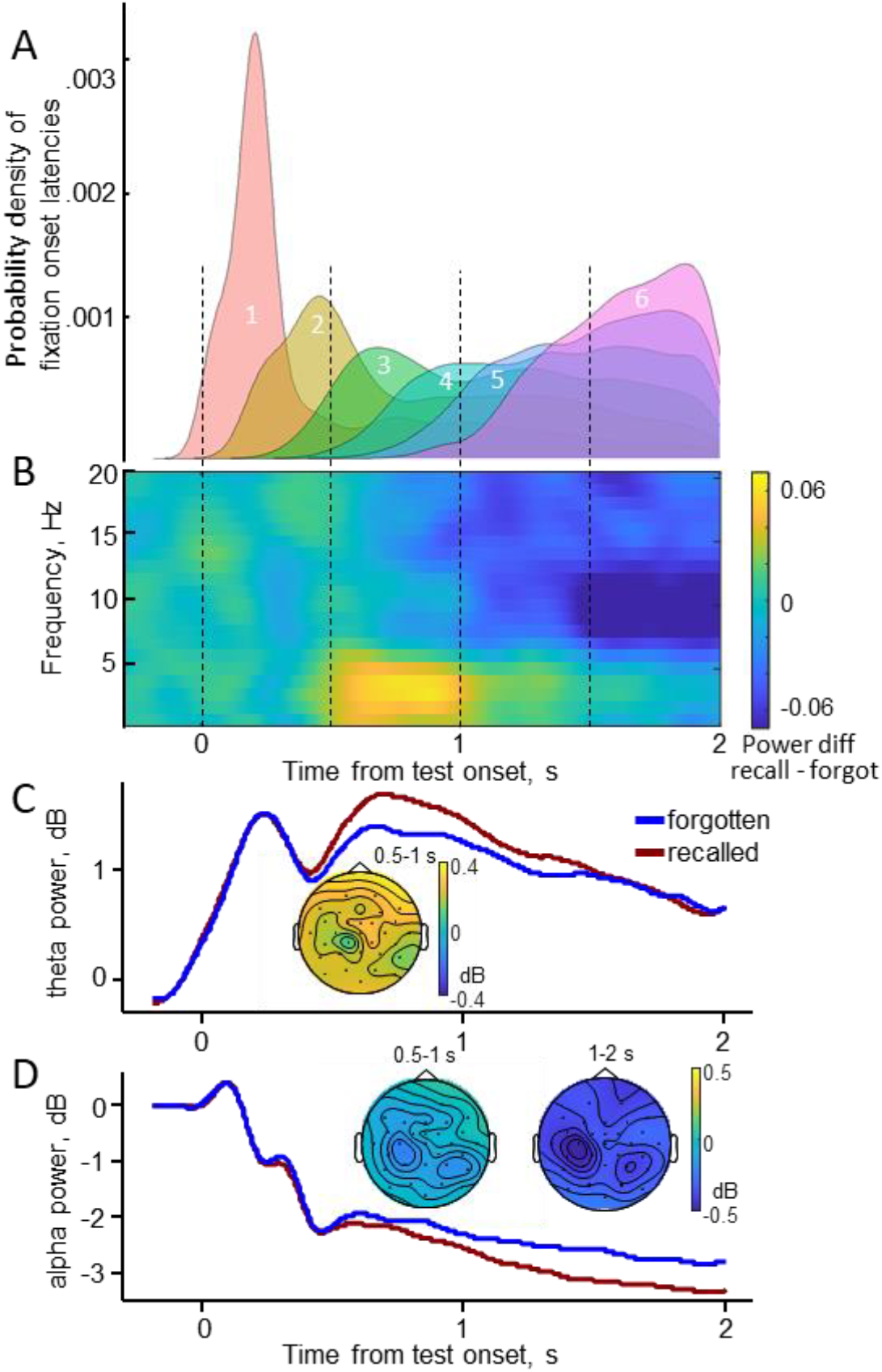
Evolution of eye movements and EEG over the entire 2-s test trial. (A) Probability density of fixation onset latencies for fixations of the first 6 ranks in the memory test trial, pooled across 28 participants. (B) Grand averaged time-frequency difference plot (recalled minus forgotten trials) averaged over all electrodes for the entire 2-s memory test trial duration. (C) theta and (D) alpha power time-locked to the test display onset for the recalled vs. forgotten trials. EEG power was baseline corrected to the −200-0 ms interval before the test screen onset, grand averaged across 28 participants and averaged over all electrodes. The insets show the topography of the memory effects as difference maps in the windows with significant differences for theta (C) and alpha (D) power.

The maximum power differences between recalled and forgotten trials in Fig. 3B and S1B are observed at 3-5 Hz in the 500-1000 ms interval and at 8-13 Hz in the 1000-2000 ms interval. Accordingly, we extracted the instantaneous power at 3-5 Hz for the theta band and at 8-13 Hz for the alpha band and averaged the power values within these early and late time windows for statistical analysis.

In the theta band, we found higher power for recalled compared with forgotten trials in the early time window (F(1, 27) = 22.4, p < .001), but no difference in the late window (Fig. 3C, S1A). We ensured that the frontal topography of this effect could not be explained by residual ocular artifacts (see the Supplementary materials for details). In the alpha band, we found lower power in the recalled than in the forgotten trials in both the early (F(1, 27) = 11.5, p = .002) and late time windows (F(1, 27) = 43.1, p < .001) (Fig. 3D, S1A).

In sum, on the time scale of the entire test trial, successful recall was predicted by higher theta power in the 500-1000 ms interval after the test screen onset and by lower alpha power over the extended 500-2000 ms interval. The higher theta combined with lower alpha power is a well-documented neural signature of successful recall in previous research where stimuli were presented at fixation (e.g., Hanslmayr et al. 2016).

### Neural activity across eye movements

To uncover the neural correlates of correct recall reflected in the relational eye-movement effect, we compared the neural correlates of recalled and forgotten trials in the EEG time-locked to the onset of fixations of first four ranks (Fig. 2B). Given that rank 1 fixations were predominantly located on the cue and fixations of subsequent ranks (2-4) were primarily located on the target and distractor (see Fig. 2A), we considered fixation rank 1 as cue-related and fixation ranks 2 through 4 as target and distractor related. Accordingly, we performed separate deconvolutions for fixation rank 1 on the cue, for fixation rank 2 on the target and distractor, and fixation ranks 3-4 on the target and distractor. A single deconvolution model included the factorial combinations of fixation elements and memory outcome (recalled vs. forgotten). The minimum number of fixations per condition per rank was set to 30. We pooled fixations from ranks 3 and 4 to reach this number. For the analysis of cue fixations of rank 1, the minimum number was reached in 23 participants, and for the analysis of target and distractor fixations of ranks 2 and 3-4, in 20 participants. The number of gaze-related EEG epochs in each condition and fixation rank is reported in the Supplementary Table 1. To analyze EEG at cue fixations of rank 1, we used a three-factor (Memory, ROI, Hemisphere) ANOVA, as described in Methods. To analyze EEG at fixations of ranks 2 and 3-4 for target and distractor fixations, we used four-factor (Memory, ROI, Hemisphere, and Element (target vs. distractor)) ANOVAs. Deconvolution and ANOVA were performed separately for theta and alpha power and for fixation ranks 2 and 3-4.

### Gaze-related theta power

For fixation rank 1 on the cue, no significant memory effect on fixation-locked theta power was observed (Fig. 4A, S2A).

**Fig. 4.**
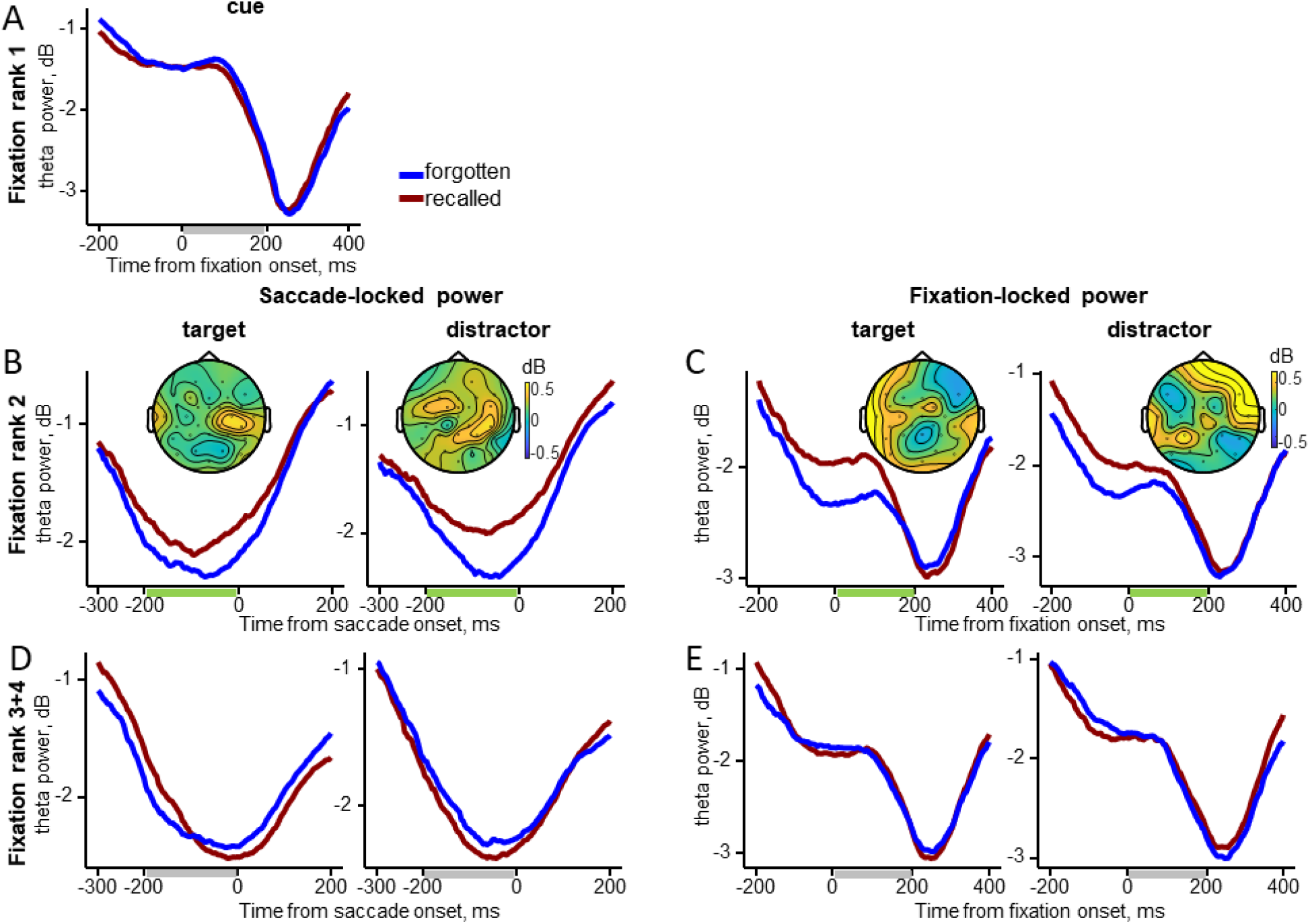
Gaze-related theta power as a function of fixation rank. Grand-averaged theta power averaged across electrodes for the recalled and forgotten trials with the difference maps (recalled minus forgotten). Time windows used for statistical analysis and mapping are shown as bars beneath the X-axis: green bars mark significant differences between recalled and forgotten trials, while gray bars mark non-significant intervals. (A) Fixation-locked power for fixations rank 1 on the cue. (B) Saccade-locked power before rank 2 fixations on the target and distractor. (C) Fixation-locked power for fixation rank 2 on the target and distractor. (D) Saccade-locked power before rank 3-4 fixations on the target and distractor. (E) Fixation-locked power for fixation ranks 3-4 on the target and distractor.

For fixation rank 2, a main effect of Memory indicated higher theta power for recalled than for forgotten trials, both in presaccadic power (F(1, 19) = 7.2, p = .015) and fixation-locked power (F(1, 19) = 13.1, p = .002) (Fig. 4B, C, S2B). No interactions emerged between Memory and Element for either time-locking types, indicating that the increase in theta power was equally pronounced for both the target and the distractor.

For fixation rank 3-4, there were no significant effects or interactions for presaccadic power. For fixation-locked power, an interaction emerged between Memory, Element and Hemisphere (F(1, 19) = 5.2, p = .03) (Fig. 4D, E, S2C, E). However, post-hoc tests revealed no significant differences (all p > .23).

Thus, for fixation rank 2, the increase in theta power associated with correct recall began during the saccade from the cue to the target or distractor and continued into the subsequent fixation. This increase did not differ statistically between target and distractor fixations. Although there was an interaction between Memory, Element and Hemisphere for fixation ranks 3-4, follow-up tests revealed no reliable differences in theta power between target and distractor. These findings suggest that the theta increase related to correct recall emerges already after cue processing, without target selectivity.

### Gaze-related alpha power

For fixation rank 1 on the cue, no significant effect of Memory on fixation-locked alpha activity was observed (Fig. 5A, S3A).

**Fig. 5.**
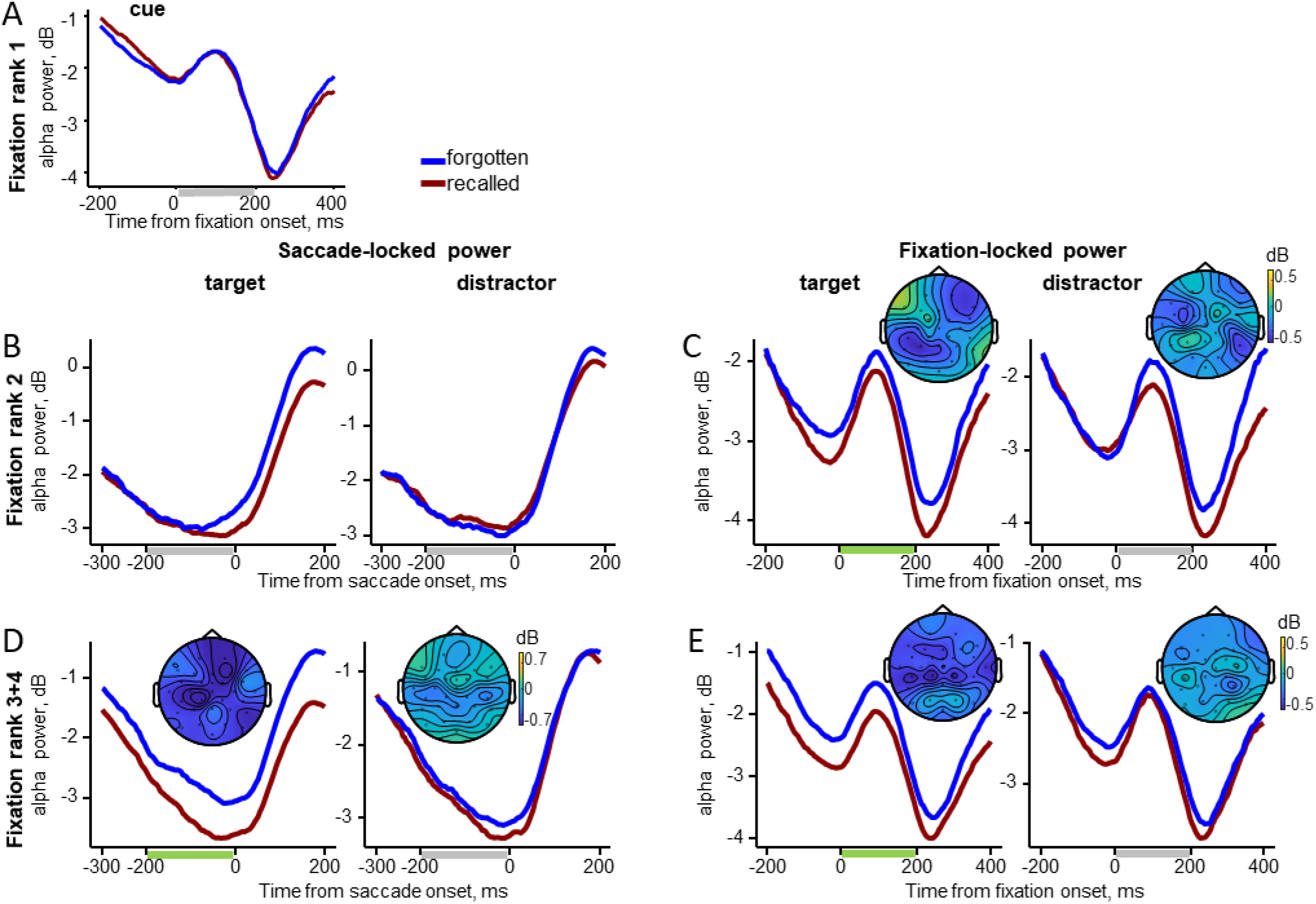
Gaze-related alpha power as a function of fixation rank. Grand-averaged alpha power averaged across electrodes for the recalled and forgotten trials with the difference maps (recalled minus forgotten). Time windows used for statistical analysis and mapping are shown as bars beneath the X-axis: green bars mark significant differences between recalled and forgotten trials, while gray bars mark non-significant intervals. (A) Fixation-locked power for fixation rank 1 on the cue. (B) Saccade-locked power before fixations rank 2 on the target and distractor. (C) Fixation-locked power for fixation rank 2 on the target and distractor. (D) Saccade-locked power before fixations rank 3-4 on the target and distractor. (E) Fixation-locked power for fixation ranks 3-4 on the target and distractor.

For fixation rank 2, for saccade-locked alpha power preceding rank 2 fixations, an interaction emerged between Memory, Element and Hemisphere (F(3, 57) = 9.0, p < .001, ε = .7). However, post hoc tests revealed no significant differences between recalled and forgotten trials (all p > .2). For fixation-locked alpha power, there was a main effect of Memory (F(1, 19) = 7.5, p = .01) indicated lower alpha power for recalled than for forgotten trials (Fig. 5C, S3B). There was an interaction between Memory, Element, ROI, and Hemisphere (F(3, 57) = 7.6, p = .002, ε = .61). Post-hoc tests indicated significantly lower alpha power in recalled compared to forgotten trials over the left parietal (p = .02) and right frontal (p = .008) areas, but only for the target (Fig. 5C, S3B, F).

To rule out the possibility that the observed interaction between Memory, Element, ROI, and Hemisphere for fixation-locked alpha power was due to participants being more accurate when targets were presented on the right side of the screen than on the left, we repeated the analysis using a dataset with an equal number of left- and right-side epochs. The results were qualitatively the same, indicating that the interaction was independent of the number of “correct” epochs on the left or right side of the screen (see the Supplementary materials for details).

For fixation ranks 3-4, the statistical results for the saccade-locked and fixation-locked alpha power were qualitatively similar (Fig. 5D, E, S3C). In both cases, there was a main effect of Memory: saccade-locked (F(1, 19) = 23.4, p < .001) and fixation-locked (F(1, 19) = 18.7, p < .001), as well as an interaction between Memory and Element: saccade-locked (F(1, 19) = 13.3, p = .002) and fixation-locked (F(1, 19) = 4.7, p = .04). Post-hoc tests showed lower alpha power for recalled than forgotten trials when fixating the target (both p < .001), but not the distractor (both p > .17).

Thus, a decrease in alpha power predictive of correct recall emerges at the second fixation, but only when the eyes land on the target. This target-selective alpha effect persists through fixation ranks 3-4 and is evident both before and after the saccade. Notably, for fixation rank 2, the decrease in fixation-locked alpha power coincides with an increase in theta power within the same time window.

In sum, the theta increase predictive of correct recall arises before the first fixation on the target or distractor, ends shortly thereafter, and does not differentiate between the two. In contrast, the alpha decrease marks successful recollection of goal-relevant associations, beginning with the first target fixation and continuing throughout the trial.

### Correlation of fixation duration with gaze-related EEG power

To examine how visual attention to the target or distractor related to gaze-related power effects, we correlated fixation duration on these elements with both saccade- and fixation-locked power. Correlations were computed separately for fixation ranks 2 and 3-4 across 20 participants, and were corrected for multiple comparisons using the false discovery rate (FDR) method. No significant correlations were observed for theta power. For alpha power, however, significant negative correlations emerged in recalled trials for the target only, indicating that longer target fixations were associated with lower alpha activity. Specifically, fixation-locked alpha power over the right occipital ROI correlated with fixation duration at rank 2 (r = −.59, p = .048), and presaccadic alpha power correlated with fixation duration at ranks 3-4 over the left frontal (r = −.60, p = .038), right frontal (r = −.56, p = .038), and parietal (r = −.54, p = .038) ROIs. These findings suggest that disproportionate target viewing (Fig. 2B) was accompanied by target-specific decreases in alpha power, particularly over the right hemisphere, consistent with the idea that greater visual attention to the target enhances the selective activation of goal-relevant associations.

## Discussion

This study investigated the neural dynamics underlying the relational eye-movement effect on a fixation-by-fixation basis during a two-alternative forced-choice associative memory test (Fig. 1). Eye-tracking results revealed a pronounced relational eye-movement effect, characterized by disproportionate viewing of targets during fixations of ranks 2-4. Next, we examined how this behavioral effect aligned with theta and alpha EEG activity, both across the entire-test trial and at the level of individual fixations.

Across the entire test trial (2000 ms), successful recall of element associations was predicted by an increase in theta power between 500-1000 ms after test screen onset and a decrease in alpha power from 500 until the end of the trial. The timing of the theta increase aligns with previous research linking theta activity to the retrieval of episodic associations (Hanslmayr & Staudigl, 2014; Herweg et al., 2020; Hsieh & Ranganath, 2014). Similarly, the timing of the alpha decrease corresponds with prior findings on its onset and evolution during episodic memory retrieval (Griffiths, Martín-Buro, Staresina, Hanslmayr, & Staudigl, 2021; Karlsson, Wehrspaun, & Sander, 2020; Martín-Buro et al., 2020; Michelmann et al., 2016).

Gaze-related EEG analyses demonstrated that the memory-related decrease in alpha power predicted retrieval of goal-relevant associations during target fixations. This effect began at the second fixation (rank 2), typically following the initial cue fixation, and unfolded gradually through fixations 2-4. In contrast, the memory-predictive theta increase was independent of gaze location: it emerged during the presaccadic interval of the second fixation; that is, before the saccade following cue fixation, regardless of whether the eyes were directed to the target or distractor, and dissipated thereafter. Together, these findings suggest that the sustained alpha effect supports the selection of relevant memory content and the reactivation of goal-relevant associations across extended fixation sequences, whereas the early, phasic theta effect predicts memory success irrespective of gaze direction, likely reflecting initial recollection of the fixated content and setting the stage for continued pattern completion of associated event elements. This temporal profile of gaze-related memory effects aligns with established stages of cued recall, which begin with cue processing within the first ∼500 ms and are followed by target reinstatement from ∼500 ms onward (Staresina & Wimber, 2019).

The observed alpha memory effect may represent an EEG correlate of recalled goal-relevant associations, as reflected in the relational eye-movement effect reported in previous research (Hannula et al., 2007; Smith et al., 2006). This interpretation is supported by the parallel time course of the alpha decrease (Fig. 5) and the disproportionate target viewing (Fig. 2B), both of which develop from the second to the fourth fixation, as well as by the negative correlation between alpha decreases and target fixation duration during correct recalls. The role of alpha, therefore, appears central to the relational eye-movement effect, consistent with prior work linking alpha desynchronization to episodic memory retrieval (Martín-Buro et al., 2020; Michelmann et al., 2016).

The relational eye-movement effect emerges quite early – well before explicit responses and sometimes even without conscious awareness (Hannula & Ranganath, 2009; Hannula et al., 2007; Nagy & Király, 2018). In line with these findings, our results suggest that processes at the second fixation of a trial following the initial cue fixation correspond to this early, potentially implicit stage of retrieval (cf., Kumaran & Wagner, 2009). Given the tight link between eye movements and attentional direction, as well as the role of attentional shifts toward a target location in signaling the timing of memory retrieval (Kragel & Voss, 2022), the disproportionate target viewing tied to the target-specific alpha memory effect likely marks the onset of associative retrieval at the second fixation.

From the second through fourth fixations, disproportionate target viewing and the alpha effect unfolded together. Prolonged visual attention to the target may facilitate activation of relevant memory content. The sustained alpha decrease may indicate the ongoing extraction of target-related information and its gradual embedding into explicit awareness (e.g., recognition of event elements) (Babiloni, Vecchio, Bultrini, Romani, & Rossini, 2006; Kumaran & Wagner, 2009; Mathewson, Gratton, Fabiani, Beck, & Ro, 2009). During retrieval tasks, decreases in alpha power have been linked to the amount of information recovered from episodic memory (Griffiths et al., 2021; Hanslmayr et al., 2016; Karlsson et al., 2020; Martín-Buro et al., 2020). The fact that this alpha reduction was observed only for the target suggests that target fixations provide more memory-relevant information than distractor fixations.

The fronto-parietal topography of the alpha effect at the second target fixation may reflect top-down control of visual attention mediated by a dorsal attentional network that includes the right frontal eye field (FEF) in prefrontal cortex and posterior cortical regions around the intraparietal sulcus (IPS) (Corbetta & Shulman, 2002). Specifically, a virtual lesion in the right prefrontal cortex during spatial attention tasks impairs target identification and reduces lateralized attention-related alpha amplitude. This finding supports a causal role for fronto-parietal alpha activity in the top-down guidance of visuospatial attention (Capotosto, Babiloni, Romani, & Corbetta, 2009; Sauseng, Feldheim, Freunberger, & Hummel, 2011). Given the involvement of right prefrontal cortical regions in episodic memory retrieval (Lepage, Ghaffar, Nyberg, & Tulving, 2000), the right-frontal alpha effect at the second target fixation likely reflects top-down control processes that facilitate selective gaze-related reactivation of target representations leading to successful recall.

Whereas alpha activity appeared tightly coupled to the selective processing of goal-relevant associations, theta activity showed a different profile. The theta power increase predicting correct recall was independent of gaze direction. This effect emerged already during the presaccadic interval of the second fixation, i.e., while the eyes were still on the cue and a saccade to the target or distractor was being planned. These findings suggest that retrieval processes are initiated during the initial cue fixation. In the present study context, this likely involves recollection of the fixated content that triggers pattern completion of information associated with a cue, an essential step for successful relational retrieval (Staresina & Wimber, 2019).

These results underscore the central role of the second fixation. Importantly, a single first fixation on the cue or any other element was never sufficient for a correct response in our two-alternative forced-choice task. Thus, the second fixation emerged as critical for accurate performance. At this stage, disproportionate target viewing co-occurs with both the early theta increase, which predicted memory success independently of whether the subsequent fixation landed on the target or distractor, and the target-specific alpha decrease. These behavioral and neural markers may reflect distinct retrieval strategies depending on the sequence in which elements are fixated.

In most trials, the first fixation was directed to the cue, followed by the target or distractor with roughly equal probability (Fig. 2A). In about 20% of trials, however, the first fixation landed on the target or distractor. This raises the possibility that participants employed different retrieval strategies depending on whether the first two fixations included the cue and target versus any combination of the cue or target with the distractor. When the cue and target were both fixated early, participants may have retrieved the task-relevant cue-target association. This dominant strategy would be consistent with the sustained, target-specific alpha effect observed from the second to fourth fixation. Alternatively, when the first two fixations included a distractor, participants may have engaged in a “recall-to-reject” strategy: recognizing the distractor and identifying that it originated from a different event than the cue or target, thereby supporting correct rejection. This strategy may be reflected in the theta effect observed during the pre- and postsaccadic intervals of the second fixation on the distractor. More broadly, these observations suggest that memory retrieval is not a unitary process but dynamically shaped by the sequence of gaze shifts. Future studies should therefore take fixation order into account to fully capture the interaction between eye movements and memory retrieval.

The primary advantage of gaze-related EEG analysis is its ability to provide a detailed, time-resolved view of the neural activity underlying the processes identified across the entire test trial analysis. For example, the onset of the alpha effect in the entire test trial aligns with the alpha effect at rank 2 (Fig. 3). This alpha effect becomes progressively stronger as the task progresses, reflected in a more pronounced alpha decrease from early to later time windows in the entire-test analysis (Fig. 3D), as well as across fixations from rank 2 to ranks 3-4 (the main effect F-values = 7.5 and 18.7, respectively, Fig. 5).

In contrast, the relationship between the gaze-related and entire test trial theta effects needs further consideration. While the theta memory effect at fixation rank 2 aligns with the onset of the theta effect observed in the entire test trial (Fig. 3C), later fixations (ranks 3-4) do not show this theta effect (Fig. 4), even though their onset falls within the theta effect interval identified in the test-level analysis (Fig. 3A). This discrepancy may arise from several factors. First, the EEG analysis of the entire test trial benefits from greater statistical power due to the higher number of trials included. Second, variations in preceding gaze sequences may obscure later fixation-related effects. For instance, fixation rank 3 on the target could be preceded by different types of fixation sequences (e.g., cue-distractor or cue-target). For fixation rank 4, the number of possible preceding sequences increases further (e.g., cue-distractor-distractor or cue-target-distractor). In some cases, retrieval-related theta activity may occur before fixation ranks 3-4, rendering the effect undetectable at those fixations. Our study was unable not isolate and analyze these variations in gaze sequences across the cue, target, and distractor elements. Future research should utilize paradigms specifically designed to track EEG activity across a variety of fixation sequences to capture these nuances and facilitate more generalizable conclusions.

Although eye movements in our paradigm followed a stereotyped pattern, they were not restricted. Analyzing EEG data coregistered with unrestricted eye movements poses specific challenges, including overlapping effects of successive saccades and condition-specific influences of eye movements (Dimigen et al., 2011; Nikolaev et al., 2016). These effects may distort, mask, or confound the effects of experimental conditions on the EEG. This issue is particularly relevant for studies of the relational eye-movement effect, as disproportionate target viewing prolongs fixation duration, making eye movements and corresponding gaze-related EEG incompatible across conditions. Addressing these challenges is critical for free-viewing EEG research. In this study, we used deconvolution modeling (Dimigen & Ehinger, 2021) to isolate the neural correlates of memory retrieval during unrestricted eye movements, providing a robust approach to mitigate these confounds.

Another key consideration in EEG-eye movement coregistration during unrestricted viewing is the selection of baseline and time windows for analysis. In traditional stimulus-response paradigms, where most information arrives at stimulus onset, EEG baselines are typically taken before the stimulus and time windows several hundred milliseconds after stimulus onset. However, in free-viewing tasks, information unfolds sequentially across fixations, and processing builds cumulatively rather than restarting with each fixation (Kragel & Voss, 2022). Consequently, fixation-related EEG may reflect not only the brain’s response to new information but also its modulation by previous representations. This sequential input poses challenges for baseline selection and time window determination (Nikolaev et al., 2016). In this study, we normalized EEG power in the continuous EEG recordings without applying traditional baseline correction. This allowed us to capture the dynamics of EEG activity across fixations. However, this approach does not account for the phase of ongoing oscillations, which is reset with each saccade (Hoffman et al., 2013; Ito, Maldonado, Singer, & Grün, 2011; Nikolaev et al., 2016). Moreover, the distinct phases of hippocampal theta oscillations precede retrieval-guided and novelty-driven eye movements, possibly coordinating switches between encoding and retrieval during free viewing (Kragel et al., 2020). In relational memory retrieval, the cycle of ongoing theta oscillations organizes sequential inputs by simultaneously representing current and previously visited items or locations (Herweg et al., 2020). Future fixation-related EEG studies of relational memory should investigate how specific phases of theta influence retrieval success to elucidate the underlying neural mechanisms further.

The specific temporal dynamics of fixations and neural activity reported in the present study should be interpreted within the context of existing research on the relational eye-movement effect. Unlike previous studies that typically employed single cue-target associations and included several distractors in their retrieval tasks (Chua et al., 2012; Hannula & Ranganath, 2009; Hannula et al., 2007; Smith & Squire, 2018; Urgolites et al., 2018), our study was designed to evaluate fixations on both target and distractor elements within scenarios featuring multiple potential cue-target associations. This methodological variation might affect the specific roles of fixations following the initial cue fixation across different paradigms. However, the general pattern of interplay between gaze behavior and neural activity in the theta and alpha bands observed in our study is likely consistent across different memory retrieval scenarios, regardless of specific paradigm parameters.

In summary, our findings underscore the active role of eye movements in relational memory retrieval. Two distinct neural signatures accompanied retrieval across successive fixations: a phasic theta increase that predicted memory success irrespective of gaze location, and a sustained, target-specific alpha decrease. Both effects were time-locked to the early fixations of each trial and co-occurred during the second fixation, when the first signs of the relational eye-movement effect emerged. At this stage, disproportionate target viewing coincided with a theta increase predictive of retrieval success and an alpha decrease reflecting the reactivation of goal-relevant associations.

These results highlight the second fixation as a critical stage in relational retrieval, where complementary neural dynamics and gaze behavior converge to support memory. Future research should build on these findings by investigating the neural and behavioral mechanisms of relational memory retrieval in more ecologically valid settings, such as tasks embedded in naturalistic environments and involving rich, contextual associations. Such investigations will deepen our understanding of how memory processes integrate with active exploration to guide goal-relevant behavior.

## Acknowledgments

We thank Axel Ekström for help with data collection and Jessica Santiago for help with initial data analysis.

## Ethics statement

Data collection was conducted in accordance with the Swedish Act Concerning the Ethical Review of Research Involving Humans (2003:460), the Code of Ethics of the World Medical Association (Declaration of Helsinki), and the local ethical guidelines at Lund University. As established by Swedish authorities and specified in the Swedish Act concerning the Ethical Review of Research involving Humans (2003:460), the present study does not require specific ethical review by the Swedish Ethical Review Authority due to the following reasons: (1) it does not deal with sensitive personal data, (2) it does not use methods that involve a physical intervention, (3) it does not use methods that pose a risk of mental or physical harm, (4) it does not study biological material taken from a living or dead human that can be traced back to that person (https://etikprovningsmyndigheten.se/wp-content/uploads/2024/05/Guide-to-the-ethical-review_webb.pdf). The data collection was anonymous and did not involve recording any potentially identifying demographic information.

## Data Availability Statement

The data that support the findings of this study are available from the corresponding author upon reasonable request.

## Credit authorship contribution statement

AN: Conceptualization, Methodology, Software, Formal analysis, Funding acquisition, Investigation, Project administration, Visualization, Writing – original draft, Writing – review & editing.

RJ: Conceptualization, Funding acquisition, Investigation, Methodology, Writing – review & editing.

IB: Conceptualization, Investigation, Methodology, Writing – review & editing.

MJ: Conceptualization, Investigation, Supervision, Writing – review & editing.

