## Supplementary materials2 for "Neural dynamics of relational memory retrieval across eye movements"

Andrey R. Nikolaev, Roger Johansson, Inês Bramão, Mikael Johansson
Lund Memory Lab, Department of Psychology, Lund University, Lund, Sweden

### Control analysis for the potential bias of memory performance by spatial congruence of images

Although locations of the encoding and test images were randomized, as described in Methods, the locations of the target and distractor images on the test screens may occasionally coincide with the locations of the respective images on the study screens. The spatial congruence between locations of the same images on the study and test screens may be recalled, which could bias memory performance. To rule out this possibility, we determined if the target appeared on the same side of the test screen relative to the vertical midline (“match”), on the opposite side (“mismatch”), or overlapped the midline (“neutral”) compared to its original encoding position. Then, for each of these three spatial-congruence conditions, we computed the number of trials, the number of correct responses, and the percentage of correct responses. The number of match and mismatch trials was about equal: 285.6 ± 19.9 and 285.1 ± 16.8 (mean ± SD), respectively, ($t = 0.1$, $p = .9$). The number of correct responses was 194.1 ± 29.7 for match and 197.4 ± 25.5 for mismatch conditions ($t = -0.7$, $p = .5$). The percentage of correct responses was 68.9 ± 8.8 for match and 69.2 ± 7.6 for mismatch conditions ($t = -1.7$, $p = .11$). So the number and percentage of correct responses did not differ between the match and mismatch conditions. In fact, the numbers were slightly higher for the mismatch, spatial-incongruent condition. This excluded the effect of spatial congruence on memory performance.

### Control analysis for the potential effect of cue image category on memory performance

The category of the cue image may affect memory performance. To control for this potential effect, we compared memory accuracy across faces, places, and objects. The percentage of correct responses was 67.5 ± 8.4 (mean ± SD) for faces, 68.4 ± 8.8 for places, and 70.0 ± 7.0 for objects. A repeated-measures ANOVA on the percentages did not reveal a significant difference across cue categories ($F(2, 38) = 2.9$, $p = .08$, $\varepsilon = .87$) (all post-hoc ps > .06). These results ruled out a systematic effect of cue image category on memory performance.

### Control analysis on the equalized left/right epoch dataset

Because the hemispheric EEG asymmetry and the related interaction between Memory, Element, ROI, and Hemisphere for fixation-locked alpha power for fixation rank 2 might have resulted from an uneven distribution of fixation-locked epochs containing targets (i.e., correct responses) on the left versus right sides of the memory test screen, we repeated all analyses on a dataset in which the number of left/right epochs was equalized. Note that any imbalance could only have occurred incidentally during the selection of epochs for the recalled and forgotten conditions, since the experiment ensured an equal number of memory tests with left/right target locations and randomized their presentation order.

Equalization was performed for fixation ranks 2 and 3-4, which included both targets and distractors. The number of left/right epochs was equalized by randomly selecting a subset from the side with more epochs to match the count from the side with fewer epochs, applied separately for targets and distractors. On average, 26% (SD = 7.5%) of the epochs were removed during this procedure. Deconvolution was then performed on the equalized datasets for theta and alpha power using the same parameters as in the main analysis. Mean power values were extracted from the 0-200 ms window following fixation onset. A repeated-measures ANOVA was conducted on these power values with the factors Memory (forgotten vs. recalled), Element (target vs. distractor), ROI (frontal, central, parietal, occipital), and Hemisphere (left vs. right).

For rank 2 fixations, a main effect of Memory ($F(1, 19) = 5.7$, $p = .03$) showed that alpha power was lower for the recalled condition than for the forgotten condition. A significant four-way interaction was observed between Memory, Element, ROI, and Hemisphere ($F(3, 57) = 6.8$, $p = .002$, $\varepsilon = .72$). Post-hoc results, however, did not reach significance: differences in the left parietal and right frontal regions for targets, which were significant in the main analysis, yielded p-values of .13 and .12, respectively. Thus, the control analysis with an equal number of left and right epochs yielded the expected decrease in statistical power but preserved the overall pattern of results from the main analysis, including the interactions involving Memory and Hemisphere. These findings rule out the possibility that the hemispheric EEG asymmetry observed in the main results was due to an incidental imbalance of target epochs on the left versus right sides of the memory test screen.

#### Supplementary Table 1

Number of fixation-related epochs per participant, the mean (SD) across 23 participants for rank 1 fixations and 20 participants for rank 2-4 fixations.

| fixations | recalled | | forgotten | |
|---|---|---|---|---|
| rank 1 - cue | 70 (23.7) | | 86 (28.9) | |
| | target | distractor | target | distractor |
| rank 2 | 59 (14.5) | 55 (11.1) | 74 (20.5) | 82 (22.1) |
| rank 3 and 4 | 86 (26.3) | 83 (27.3) | 104 (42.1) | 129 (48.6) |

#### Supplementary figures

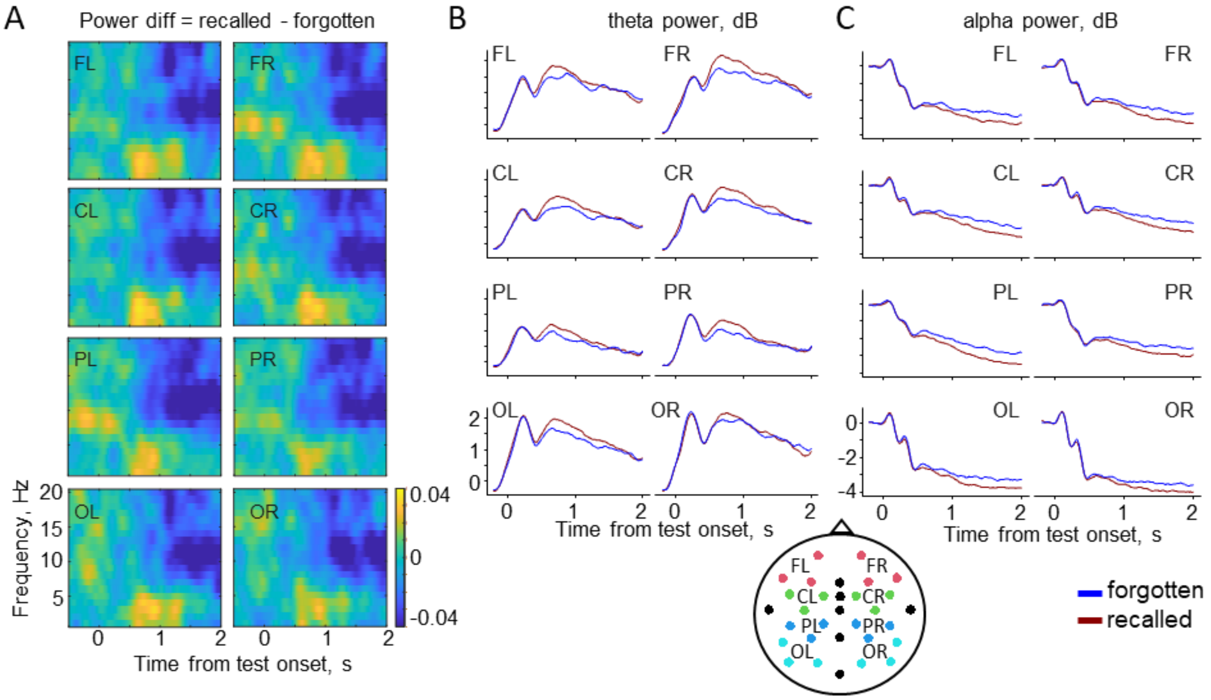


Fig. S1. **EEG results for the entire duration of the memory test trial.** (A) Time-frequency difference plots (recalled minus forgotten conditions) for 8 ROIs: Frontal, Central, Parietal and Occipital brain regions of the Left and Right hemispheres (FL, FR, CL, CR, PL, PR, OL, OR). Theta (B) and alpha (C) power time-locked to the test display onset for the forgotten vs. recalled conditions. Power was baseline corrected to the -200-0 ms interval before the test screen onset and grand averaged across 28 participants and across the triples of electrodes shown in the inset below the panels, resulting in 8 ROIs.

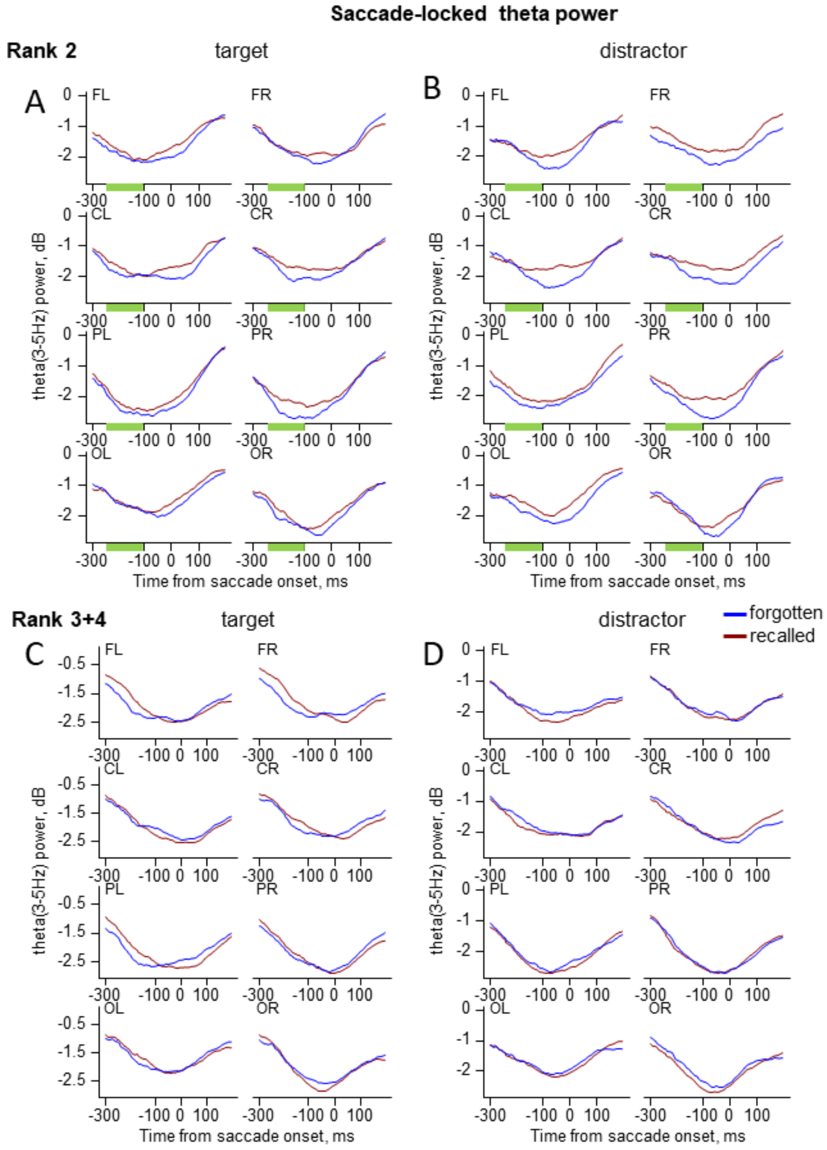


Fig. S2. **Saccade-locked theta power as a function of fixation rank**. (A-D) The theta power grand-averaged across participants in 8 ROIs for cue, target, and distractor for the forgotten vs. recalled conditions. (A) Presaccadic power before rank 2 fixations on the target; (B) Presaccadic power before rank 2 fixations on the distractor; (C) Presaccadic power before rank 3 and 4 fixations on the target; (D) Presaccadic power before rank 3 and 4 fixations on the distractor. Significant differences between for the forgotten and recalled conditions are indicated by green bars.

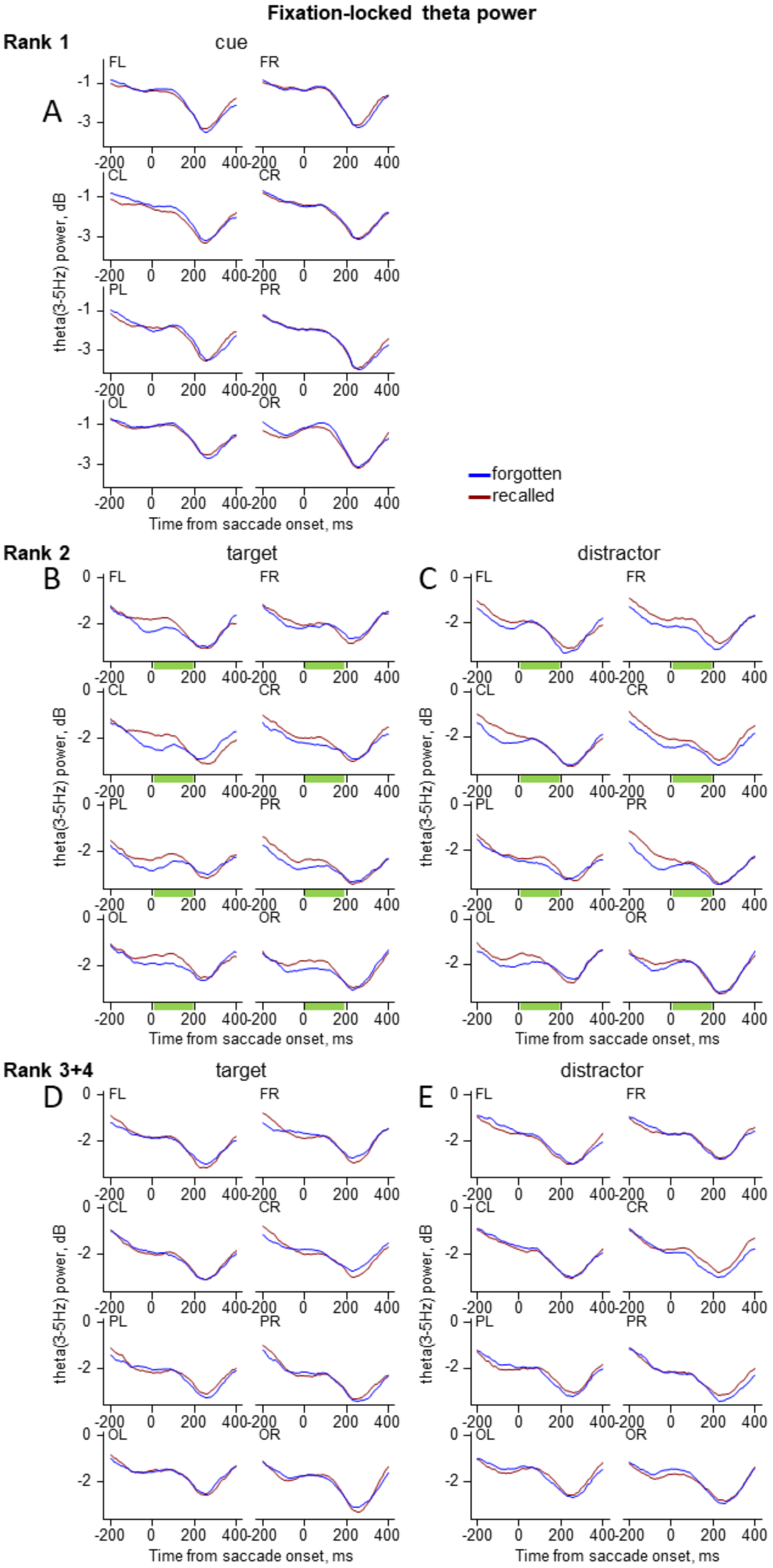


Fig. S3. **Fixation-locked theta power as a function of fixation rank**. (A-E) The theta power grand-averaged across participants in 8 ROIs for cue, target, and distractor for the forgotten vs. recalled conditions. (A) Theta power for rank 1 fixations on the cue; (B) Theta power for rank 2 fixations on the target; (C) Theta power for rank 3 and 4 fixations on the target; (D) Theta power for rank 2 fixations on the distractor; (E) Theta power for rank 3 and 4 fixations on the distractor. Significant differences between for the forgotten and recalled conditions are indicated by green bars.

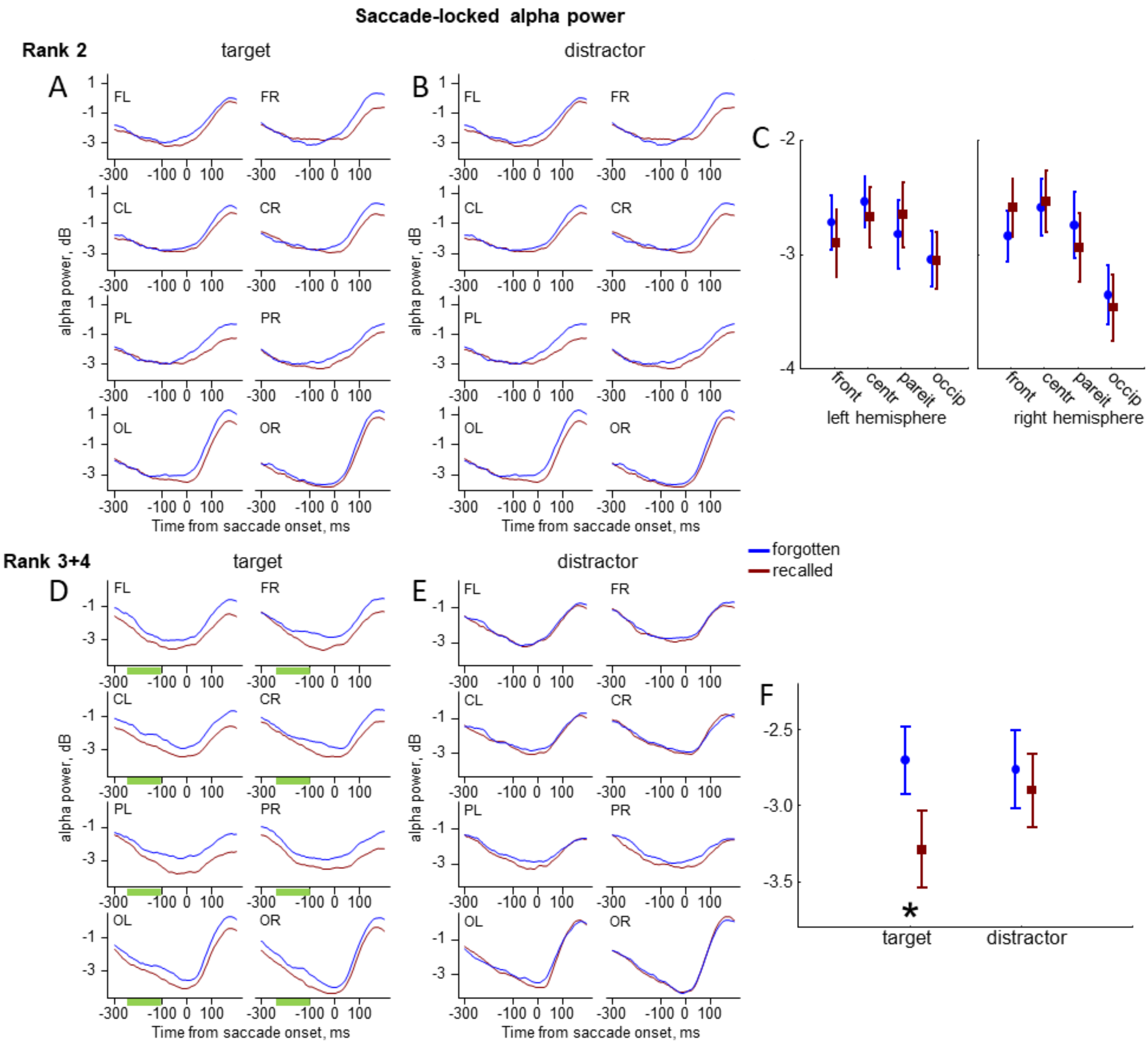


Fig. S4. **Saccade-locked alpha power as a function of fixation rank.** (A, B, D,E) The grand averaged alpha power across participants in 8 ROIs for cue, target, and distractor for the forgotten vs. recalled conditions. (A) Presaccadic power before rank 2 fixations on the target; (B) Presaccadic power before rank 2 fixations on the distractor; (C) A mean-error plot of the interaction between Memory, ROI, and Hemisphere in the -200-0 ms interval before the onset of the saccade that leads to rank 2 fixations. Error bars indicate standard errors of the mean across participants. (D) Presaccadic power before rank 3 and 4 fixations on the target; (D) Presaccadic power before rank 3 and 4 fixations on the distractor. Significant differences between forgotten and recalled conditions are indicated by green bars. (F) A mean-error plot of the interaction between Memory and Element in the -200-0 ms interval before the onset of the saccade that leads to rank 3-4 fixations. Asterisks indicate significant differences between the memory conditions.

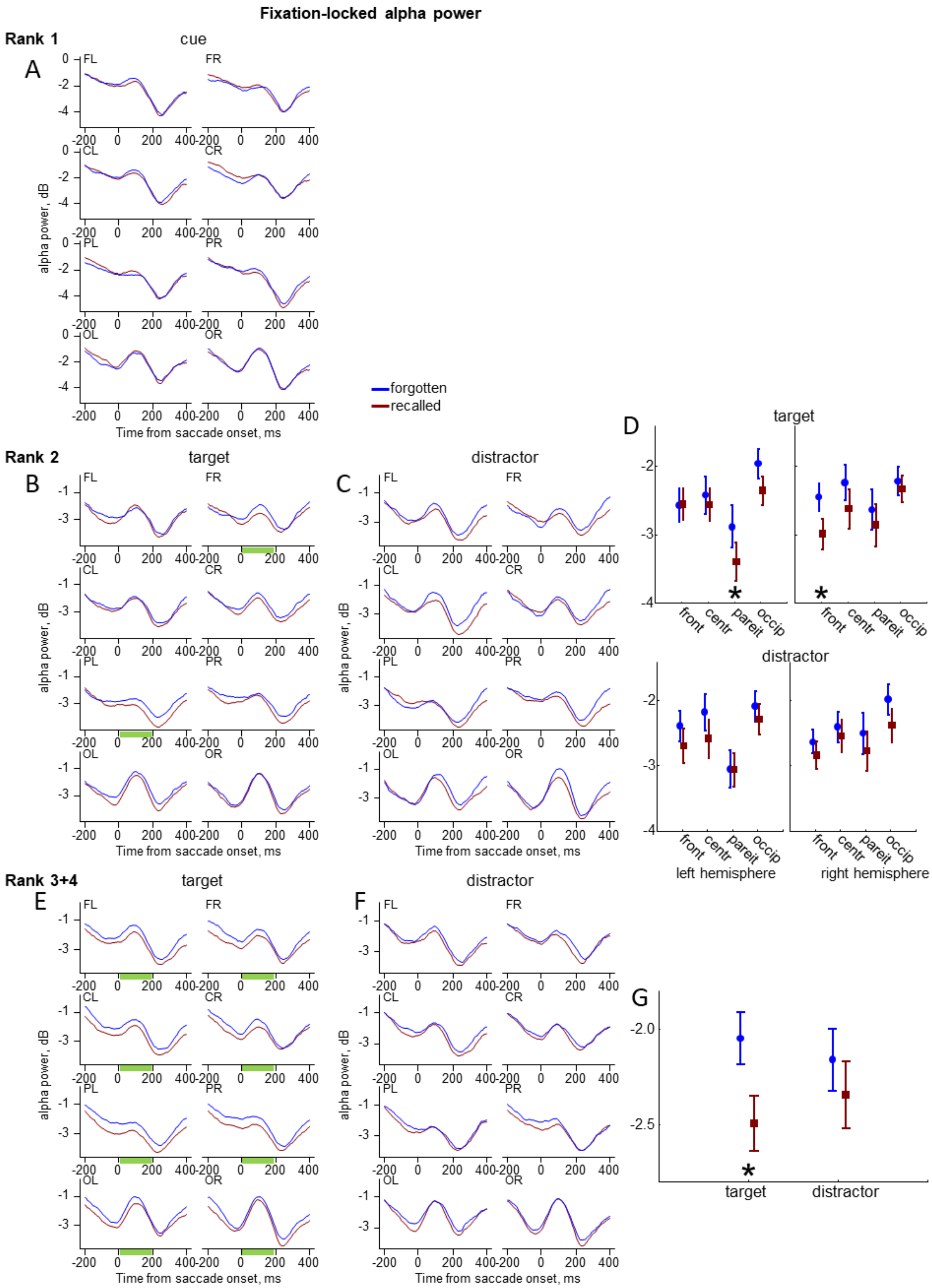


Fig. S5. **Fixation-related alpha power as a function of fixation rank**. (A-C, E, F) The grand averaged alpha power across participants in 8 ROIs for cue, target, and distractor for the forgotten vs. recalled conditions. (A) Alpha power for rank 1 fixations on the cue; (B) Alpha power for rank 2 fixations on the target; (C) Alpha power for rank 2 fixations on the distractor; (D) A mean-error plot of the interaction between Memory, Element, ROI, and Hemisphere in the 0-200 ms interval after the onset for rank 2 fixations. (E) Alpha power for rank 3 and 4 fixations on the target; (F) Alpha power for rank 3 and 4 fixations on the distractor. Significant differences between forgotten and recalled conditions are indicated by green bars. (G) A mean-error plot of the interaction between Memory and Element in the 0-200 ms interval after the onset for rank 3-4 fixations. Error bars indicate standard errors of the mean across participants. Asterisks indicate significant differences between the memory conditions.

#### Control for residual ocular artifacts

The frontal theta activity can be contaminated by eye movement artifacts during the 2-s memory test trial. This issue is particularly relevant in EEG-eye movement coregistration studies involving unrestricted gaze behavior (Dimigen et al., 2011; Nikolaev et al., 2016). To address this, we used a dedicated preprocessing approach optimized for free viewing: the ICA-based OPTICAT toolbox (Dimigen, 2020), which identifies and eliminates all major ocular artifacts, such as blinks, eyeball rotations, and saccadic spike potentials (see Methods for details). Due to this rigorous preprocessing, it is unlikely that the frontal theta memory effect observed in the difference maps (recalled minus forgotten) during the 500-1000 ms interval is driven by residual ocular artifacts.

Nevertheless, to rule out ocular contamination more conclusively, we considered the broadband nature of ocular artifacts (Keren et al. 2010; Plöchl et al. 2012). Specifically, if present, such broadband activity should be visible across frequency bands in the time-frequency plots for the eight ROIs (Fig. S1A), but this was not the case. Furthermore, we examined the ERPs within a frequency range that includes a major portion of ocular artifacts by filtering the EEG between 0.1 and 30 Hz. Fig. S6 presents ERPs in 8 ROIs, ERP topoplots, and a series of ERP maps across ten 200-ms intervals. They demonstrate the absence of abnormal increases in frontal activity, which would suggest ocular contamination. Since the suspicious frontal increase was observed on the differential maps (recalled minus forgotten), we next compared ERPs using the same ANOVA design as before in the 500-1000 ms interval, where the theta memory effect is found. We found no evidence of a Memory effect or its interactions with ROIs (all Fs < 0.8, all ps > .54).

Together, these observations indicate that the theta memory effect within the 500–1000 ms interval reflects genuine differences between memory conditions rather than residual ocular artifacts.

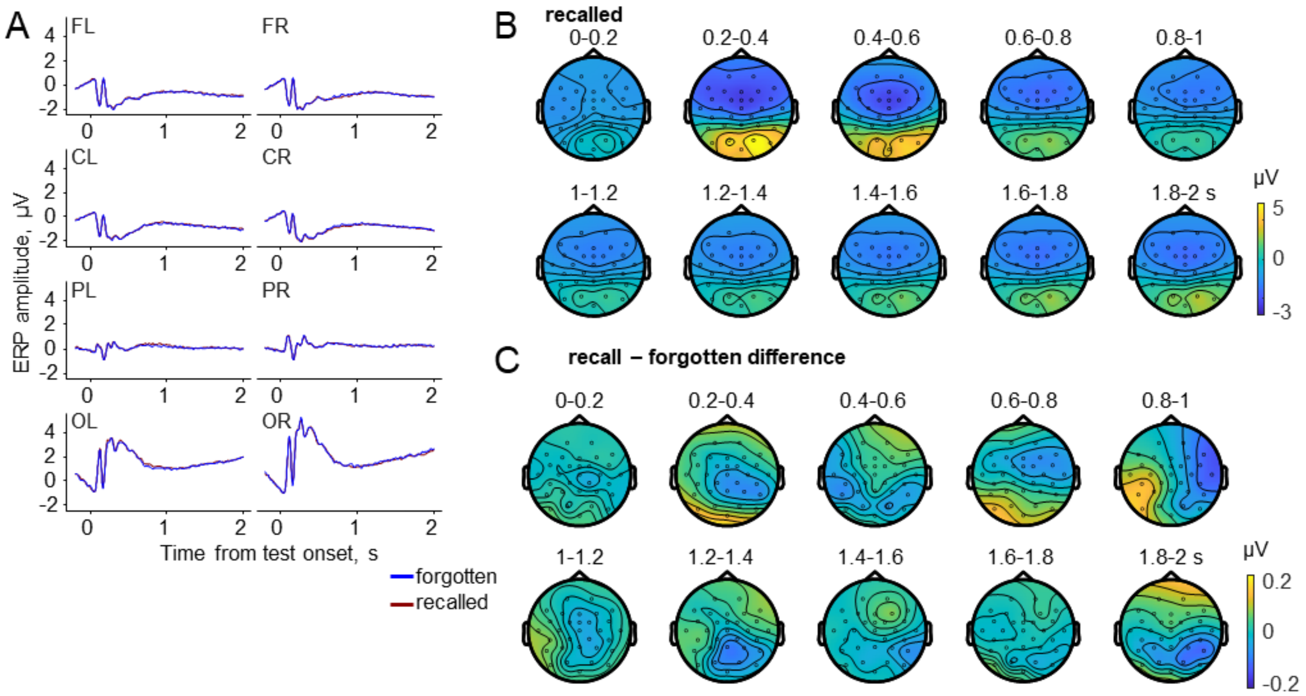


Fig. S6. **Grand-averaged (N=28) ERP filtered between 0.1 and 30 Hz for the entire 2-s memory test trial.** (A). ERP time-locked to the test screen onset for the recalled vs. forgotten trials (the difference between these conditions is hardly visible). (B) ERP maps of the means in 200 ms intervals for the recalled condition (the maps for the forgotten condition look almost the same). (C) ERP maps of the mean differences (recalled minus forgotten condition) in 200 ms intervals.

Keren, A. S., Yuval-Greenberg, S., & Deouell, L. Y. (2010). Saccadic spike potentials in gamma-band EEG: characterization, detection and suppression. Neuroimage, 49(3), 2248-2263.
Plöchl, M., Ossandón, J. P., & König, P. (2012). Combining EEG and eye tracking: identification, characterization, and correction of eye movement artifacts in electroencephalographic data. Frontiers in human neuroscience, 6, 278.
